# The ferroptosis mediator ACSL4 fails to prevent disease progression in mouse models of MASLD

**DOI:** 10.1101/2024.06.11.598353

**Authors:** Carolin Angendohr, Christiane Koppe, Anne T Schneider, Leonie Keysberg, Michael T Singer, Matthias A Dille, Johannes G Bode, Marcus Conrad, Mihael Vucur, Tom Luedde

**Affiliations:** Department of Gastroenterology, Hepatology and Infectious Diseases, University Hospital Duesseldorf, Medical Faculty at Heinrich Heine University Duesseldorf, Duesseldorf, Germany; Institute of Developmental Genetics, Helmholtz Zentrum München, Neuherberg, Germany

**Keywords:** Ferroptosis, ACSL4, MASLD, liver cirrhosis

## Abstract

Metabolic dysfunction-associated steatotic liver disease is an increasingly prevalent condition, representing a major risk factor for the development of progressive chronic liver damage. This can potentially give rise to the development of steatohepatitis and hepatocellular carcinoma (HCC). It is already known, that patients with metabolic dysfunction-associated steatotic liver disease (MASLD) show increased systemic and hepatic iron concentrations as well as perturbed lipid metabolism, suggesting involvement of ferroptotic cell death in the development and progression of MASLD. Consequently, inhibition of ferroptosis represents a potential therapeutic option for patients with MASLD.

The aim of this study was to determine whether a liver parenchymal cell specific conditional deletion of the pro-ferroptotic gene acyl-CoA synthetase long-chain family member 4 (ACSL4^LPC-KO^) can ameliorate the onset and progression of MASLD in mice. To this end, ACSL4^LPC-KO^ and floxed wild-type littermates were fed a choline-deficient high-fat diet over the course of 20 and 40 weeks to monitor the progression of metabolic liver injury as well as the development of metabolic syndrome.

In contrast to the recently published studies by Duan et al. [1], our results show no significant differences between ACSL4^LPC-KO^ and WT mice with regard to the development of MASLD or the progression of metabolic syndrome. Furthermore, no differences were observed in metabolic parameters (i.e. weight gain, glucose tolerance test, hepatic steatosis) or MASLD-associated inflammatory response. Our analyses therefore suggest that loss of ACSL4 has no effect on the progression of MASLD induced by choline-deficient high fat diet. The discrepancy between our and previously published results could be due to differences in the diets or the influence of a distinct microbiome, so the results obtained with liver parenchymal cell specific ACSL4 null mice should be taken with caution.

## Introduction

Obesity and its associated complications represent a growing global public health challenge. As such, the incidence of metabolic dysfunction-associated steatotic liver disease (MASLD) as a hepatic manifestation of metabolic disease has increased in recent years [2,3]. MASLD is defined as increased hepatic steatosis, whether due to increased alcohol consumption or dietary fat intake. An increase in steatosis can result in liver damage and inflammation, which is known as metabolic-dysfunction associated steatohepatitis (MASH). If left untreated, this can progress to cirrhosis and HCC [4]. The therapeutic options available for these patients are severely limited. Only a minority of patients are able to achieve lifestyle modification with consequent loss of body weight [5]. In contrast, bariatric surgery is highly invasive and associated with complications [6]. Consequently, there is an urgent need for non-invasive therapeutic approaches. Despite considerable efforts that have been made in metabolic science in the past years, no drug has been approved for use in patients with MASLD. Programmed cell death (PCD) has previously been linked to the pathogenesis of MASLD/MASH. However, preclinical studies on the both best studied PCD forms apoptosis [7,8] and necroptosis [9,10] showed contradictory results. Ferroptosis, a form of regulated necrotic cell death driven by iron-dependent lipid peroxidation, has been implicated in the pathogenesis of MASLD [11].

The susceptibility of cells to ferroptosis is, among others, influenced by intracellular iron levels [12] and the composition of the lipid membrane, with high levels of polyunsaturated fatty acids (PUFAs) in phospholipids promoting ferroptosis [13,14]. Patients with MASLD or MASH display elevated serum and liver iron levels, indicative of an increased ferroptosis sensitivity [15]. Furthermore, acyl-CoA synthetase long chain member 4 (ACSL4), which may contribute to ferroptosis susceptibility by preferably activating long chain PUFAs that can then be esterified into phospholipids of cellular membranes [13], is upregulated in patients with hepatic steatosis [16]. Based on this, it has recently been shown that the inhibition of ferroptosis by liver parenchymal cell specific knockout of ACSL4 in mice protects against developing obesity and associated steatosis when fed a high-fat/high-cholesterol/high-fructose diet (42% kcal fat, 42% kcal cholesterol with drinking water containing sucrose and fructose), a methionine-choline deficient diet (MCD), or a high-fat diet (60% kcal fat) [1].

In the present study, we set out to investigate the effects of a conditional liver parenchymal cell (LPC) specific ACSL4 knock out (ACSL4 ^LPC-KO^) in mice fed with a choline-deficient, high-fat diet (CD-HFD). This diet is known to efficiently recapitulate the key features of human metabolic syndrome, which encompasses weight gain and development of glucose intolerance [17]. Notably, no differences were observed between ACSL4^LPC-KO^ and WT mice with regard to the development of MASLD or metabolic syndrome. Our findings therefore challenge the prevailing view on the involvement of ferroptosis in the development of MASLD and underscore the necessity of employing diverse dietary regimens in the quest for innovative therapeutic approaches.

## Results

*ACSL4* ^LPC-KO^ *fails to provide protection against the development of metabolic syndrome on CD-HFD –* To study the effect of ferroptosis inhibition in MASLD, we deleted the pro-ferroptotic regulator ACSL4 specifically in liver parenchymal cells (LPC-KO, ACSL4^LPC-KO^) (Fig. 1a). ACSL4^LPC-KO^ and floxed wild-type littermates (WT) were fed with CD-HFD for either 20 or 40 weeks. CD-HFD feeding resulted in weight gain over the 20-week and 40-week feeding periods, however, no differences were observed between ACSL4^LPC-KO^ and WT mice in terms of relative or absolute weight gain (Fig. 1b and 1c). This was also accompanied by a similar weekly feed intake (Fig. 1d).

**Figure 1:**
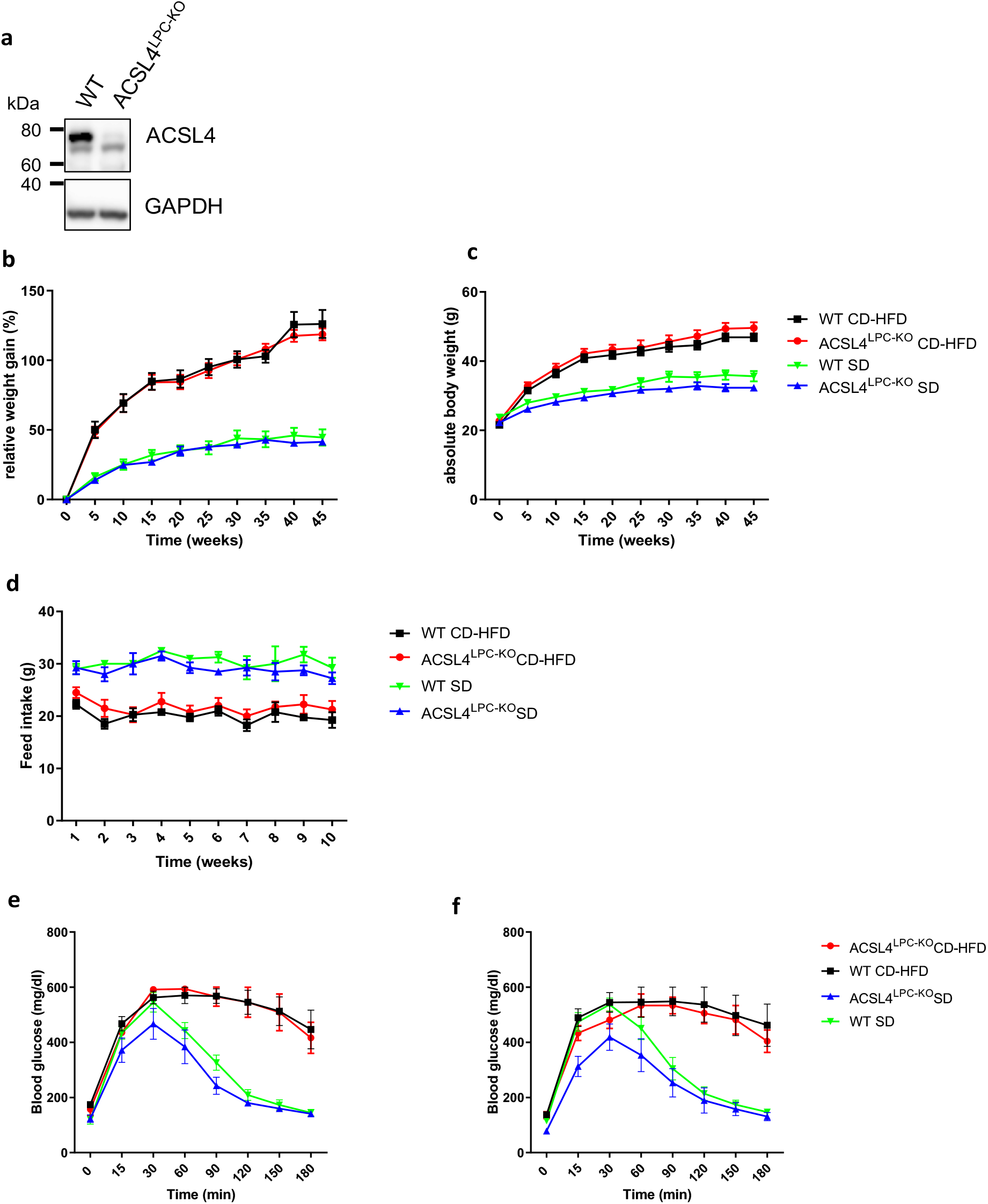
ACSL4 ^LPC-KO^ has no impact on weight gain and glucose tolerance under CD-HFD. (a) ACSL4 protein levels in the livers of ACSL4^LPC-KO^ and loxP-flanked control mice fed normal chow were assessed by Western blot. (b-c) ACSL4 ^LPC-KO^ showed similar weight gain compared to WT under CD-HFD over the indicated time period, both in terms of relative weight gain (b) and absolute weight (c). (d) Feed intake was lower in CD-HFD than in SD, but did not differ between ACSL4 ^LPC-KO^ and WT. (e-f) CD-HFD led to impaired glucose tolerance compared to SD measured by glucose tolerance tests. In 20 weeks (e) and 40 weeks old mice (f) fed with indicated diets, no difference between ACSL4 ^LPC-KO^ and WT was observed.

Since obesity and MASLD is often accompanied by the development of type II diabetes, we next examined the impact of ACSL4 ^LPC-KO^ in glucose tolerance test under CD-HFD. CD-HFD resulted in impaired glucose tolerance compared to mice fed a standard diet. In accordance with the previously observed outcomes, the deletion of ACSL4 did not improve glucose tolerance in either the 20-week feeding group (Fig. 1e) or the 40 week feeding group (Fig. 1f). To summarize, liver-parenchymal cell knockout does not protect against diet-induced weight gain and its consequences.

*ACSL4* ^LPC-KO^ *does not protect mice from developing MASLD and hepatic injury under CD-HFD –* As described above, CD-HFD not only leads to weight gain and the development of metabolic syndrome, but also to MASLD with subsequent liver damage. To investigate the influence of liver-parenchymal cell specific knockout on the development of MASLD with progression to MASH, we further characterized the hepatic phenotype of mice under CD-HFD. Macroscopic (Fig. 2a) and histopathological (Fig. 2a) examination of the liver showed pronounced steatosis, but no differences between ACSL4 ^LPC-KO^ and WT mice after 20 and 40 weeks of CD-HFD were observed. In line, the nonalcoholic fatty liver disease activity score (NAS score) also demonstrated no differences between the two groups (Fig. 2b). Furthermore, serum cholesterol and triglyceride levels were found to be similar in WT and ACSL4 ^LPC-KO^ mice after 20 (Fig. 2c) and 40 weeks of feeding with SD or CD-HFD (Fig. 2d). Since the progression of MASLD to MASH is associated with liver damage, we next measured the serum parameters for liver injury (AST, ALT, GLDH, LDH) in mice fed with CD-HFD and standard diet (SD). While CD-HFD was associated with a mild hepatitis, there were also no differences between ACSL4 ^LPC-KO^ and WT mice in the group of 20 weeks (Fig. 2c) and 40 weeks of feeding (Fig. 2d).

**Figure 2:**
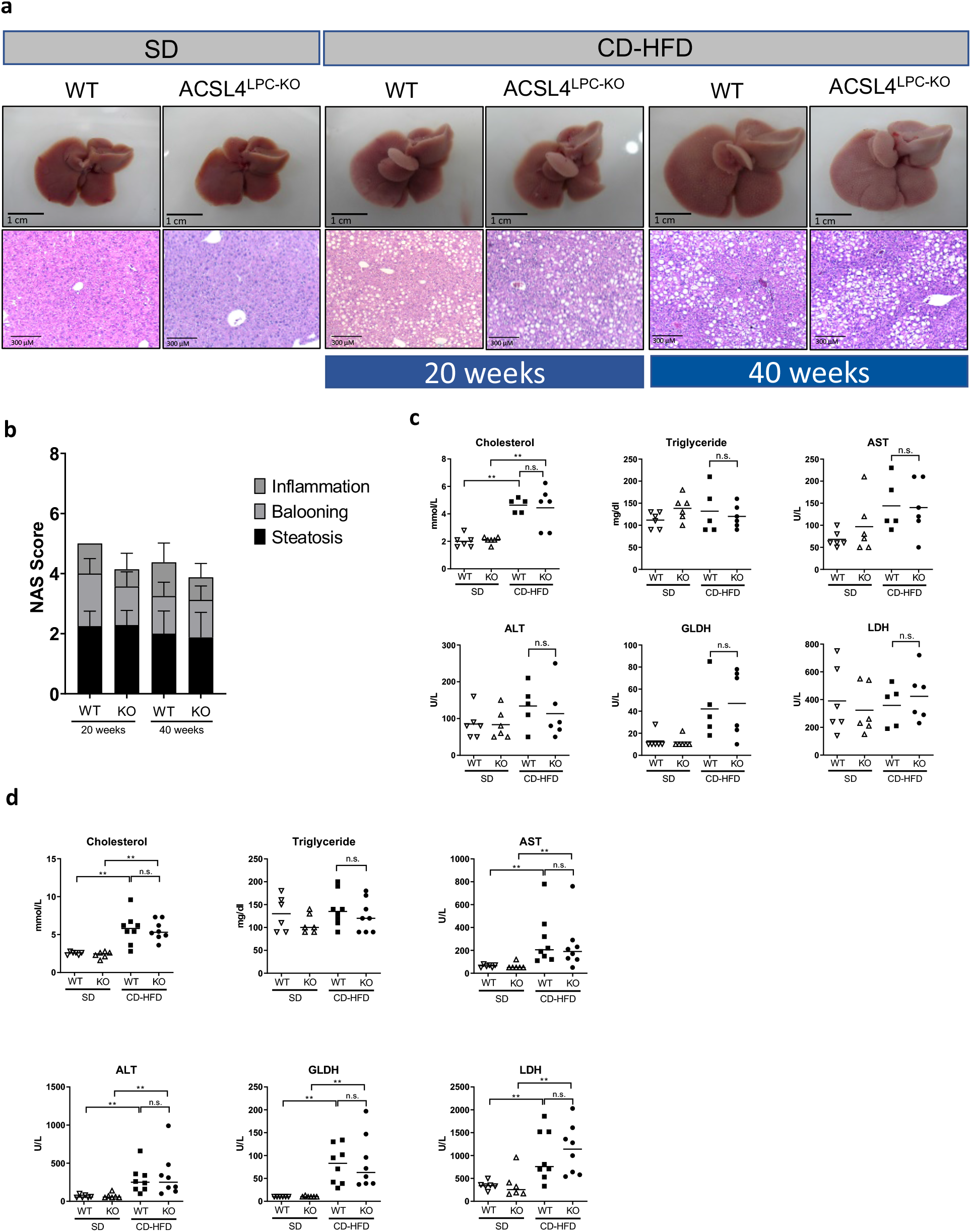
ACSL4 ^LPC-KO^ and WT mice both develop MASLD and associated liver injury after 20 and 40 weeks of feeding on CD-HFD to the same extent. (a) ACSL4 ^LPC-KO^ mice and wild type mice both develop steatosis hepatitis after 20 and 40 weeks of feeding. (b) No difference between ACSL4 ^LPC-KO^ and WT in the development of steatohepatitis was observed in the NAS Score on histology in 20 and 40 weeks old mice. (c-d) After 20 weeks (c) and 40 weeks (d), the CD-HFD led to an increase in serum values showing liver damage (AST, ALT, GLDH and LDH) and aberrant parameters of lipid metabolism (triglycerides, cholesterol) as indicated. No difference could be detected between ACSL4 ^LPC-KO^ and WT mice.

In aggregate, both ACSL4 ^LPC-KO^ and WT mice developed diet-induced liver damage as defined by MASLD or MASH to the same extent.

*ACSL4* ^LPC-KO^ *mitigates inflammation and fibrogenesis induced by CD-HFD –* As the progression of MASLD to MASH in humans is accompanied by intrahepatic inflammation with an accumulation of immune cells, we further sought to characterize inflammation upon CD-HFD feeding. It has already been described that the infiltration of different immune cells, particularly lymphocytes, contributes to the phenotype observed in CD-HFD [17]. Of note, we could not detect differences between ACSL4 ^LPC-KO^ and WT mice regarding the amount of infiltrating CD3^+^ T-lymphocytes, B220^+^ B-lymphocytes and F4/80^+^ macrophages (Fig. 3a-c). We further examined expression of the indicated cytokines and alarmins in the whole liver tissue of mice fed with CD-HFD. Again, we did not find differences in hepatic inflammation induced by CD-HFD between ACSL4 ^LPC-KO^ and WT mice (Fig. 3d).

**Figure 3:**
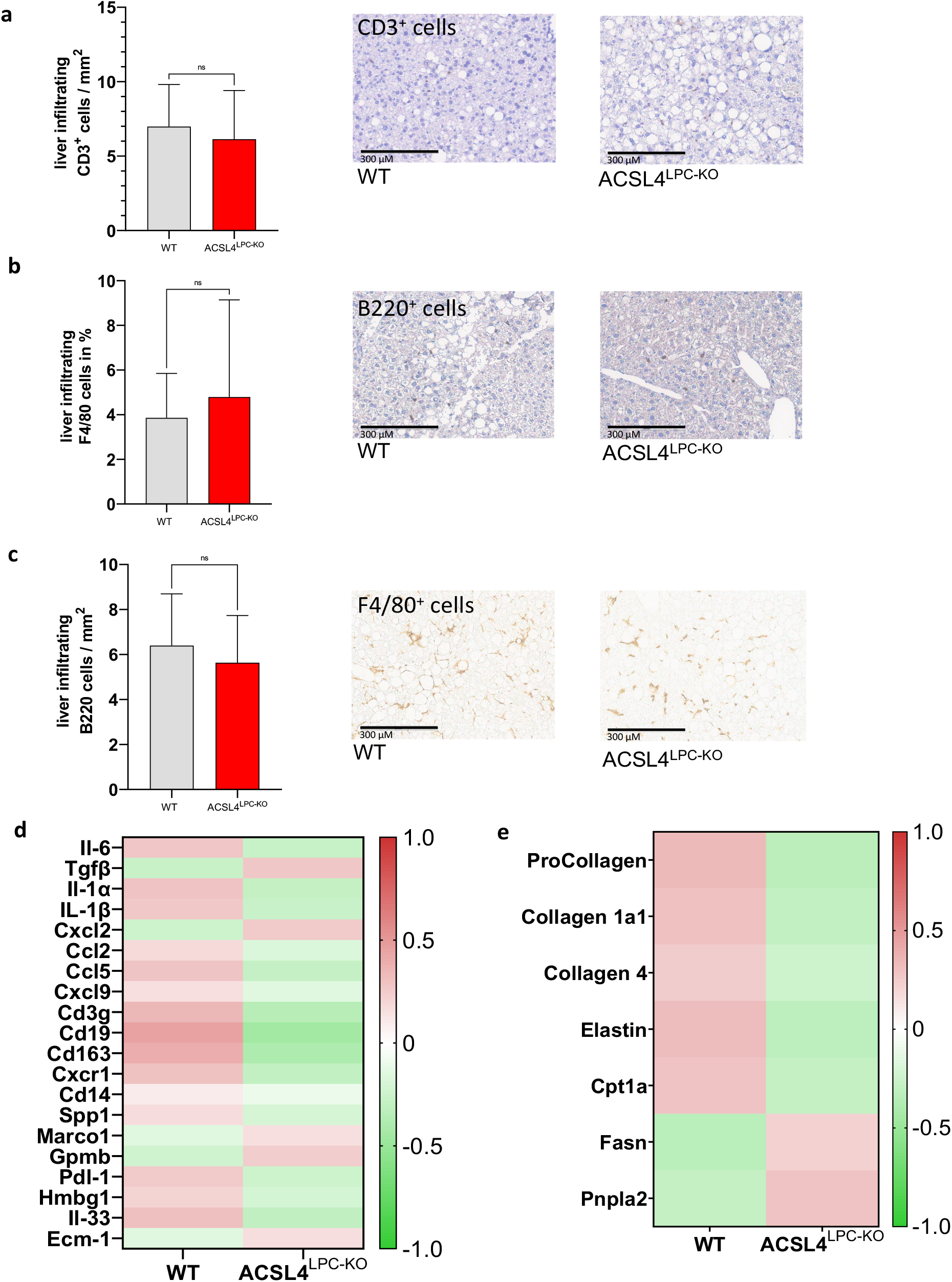
qPCR-based and immunhistochemical examinations showed no differences between ACSL4 ^LPC-KO^ and WT in terms of tissue inflammation and fibrogenesis. (a-c) Immunhistochemical staining did not show any significant differences of CD3^+^ T-cells (a) and B220^+^ B cells (b) and F4/80^+^ cells (c) between liver of ACSL4 ^LPC-KO^ and WT mice fed with a CD-HFD for 40 weeks. (d) qPCR-based studies indicated no differences in inflammatory parameters (cytokines, distinct receptors and alarmins) between ACSL4 ^LPC-KO^ and WT mice fed with CD-HFD for 40 weeks. (e) Analysis of distinct fibrogenesis and fat metabolism associated genes in whole liver tissue after 40 weeks of CD-HFD showed no difference between WT and ACSL4 ^LPC-KO^.

We finally tested expression of fibrosis-associated genes via qPCR and again we could show that genes promoting hepatic fibrogenesis and hepatic lipid deposition were not significantly downregulated in ACSL4 ^LPC-KO^ compared to WT mice (Fig. 3e).

Taken together, these findings suggest that ACSL4 deletion has no impact on the inflammatory response and fibrosis in the progression of MASLD and MASH.

## Discussion

Ferroptosis is characterized by an iron-dependent, unrestrained generation of lipid hydroperoxides in cellular membranes, which subsequently leads to membrane deterioaration, membrane rupture and cell death [18]. ACSL4 may increase the cell’s susceptibility to ferroptosis, as it preferably activates long-chain PUFAs (namely arachidonic acid and adrenic acid) which upon esterification into phospholipids by another class of enzymes may undergo peroxidation, thereby contributing to phospholipid autooxidation and cell membrane rupture [13]. Interestingly, arachidonic acid has been demonstrated to be upregulated in mice fed a MCD suggesting therapeutic potential in modulating ferroptosis in MASLD [19].

Indeed, pharmacological ferroptosis inhibitors (e.g. ferrostatin-1, liproxstatin-1 or Trolox) mitigate MASLD and inflammation associated MASH progression in mice [19-22]. Moreover, rosiglitazone, a PPARg agonist and approved drug for the treatment of diabetes mellitus, exerts its effect by inhibiting ACSL4 (as an off-target effect) and ferroptosis, leading to weight loss, and has been described to ameliorate arsenic-induced steatohepatitis in mice [23-26]. Examining the distinct role of ACSL4 in progression of MASLD, Duan et al. challenged ACSL4 ^LPC-KO^ and WT mice with three different diets to induce development of MASLD and metabolic syndrome: (i) high-fat, high-cholesterol, and high-fructose diet, (ii) methionine-choline deficient diet and (iii) high-fat diet. They found that ACSL4^LPC-KO^ mice show increased mitochondrial respiration and β-oxidation with increased catabolic fatty acid metabolism which protected against MASLD and metabolic syndrome [1]. In this work, we challenged the role ACSL4 in the same murine model of ACSL4^LPC-KO^ when subjected to a CD-HFD. CD-HFD induces steatosis, inflammation, mild fibrosis and weight gain when fed over 10 weeks [27].

Unexpectedly, we did not observe a protective effect of ACSL4^LPC-KO^ on the development of MASLD or weight gain. Moreover, there were no differences in hepatic injury, cholesterol metabolism or glucose tolerance. These contrasting results raise the question of the underlying causes.

It could be suggested that choline-deficient diet limits the synthesis of phospatidylcholine, which is a driver of ferroptosis, and thus reduces the protective effect of ACSL4^LPC-KO^. However, it has been already shown that dietary choline deficiency does not lead to a decrease in PC in mice [28]. It should be noted that investigating ferroptosis in a model of CD-HFD may present certain challenges. Previous studies have shown that deferoxamine, a common ferroptosis inhibitor, was unable to mitigate liver injury in mice fed a CD-HFD and ethanol [19]. In contrast, two other ferroptosis inhibitors were found to inhibit cell death and inflammation in the same dietary model: Trolox, a derivative of vitamin E, and deferiprone, an iron chelator [19].

Of note, the observed weight gain in the CD-HFD exceeded the weight gain observed in the work of Duan et al. when fed high-fat diet (HFD) and HFD with added sucrose and fructose in the drinking water, although the fat content was comparable or even less (60% and 42% vs. 45% for CD-HFD) [1]. The reasons for these contradictory results of our work and that of Duan et al. remain vague. The absence of choline per se does not result in an increase in body weight [29]. However, variations in genetic background or a possible influence by the microbiome or husbandry conditions throughout the different animal facilities may reason these disparate phenotypes [30,31].

The progression from MASLD to MASH, cirrhosis and eventually HCC in human and mice is triggered by increased intrahepatic inflammation [17,32]. In this context, it is widely acknowledged that ferroptosis as a lytic form of cell death results in the release of intracellular components, which in turn may trigger immune activation and remodel the immunological microenvironment [33,34]. Given that ferroptosis inhibitors have been demonstrated to alleviate inflammatory damage in mice [20], we hypothesized that ACSL4^LPC-KO^ should result in an attenuation of hepatic inflammation and subsequent fibrogenesis. As lymphocytes and macrophages are the primary cells involved in the pathogenesis of MASH [32,35], we examined the amount of infiltrating B- and T-lymphocytes and F4/80 macrophages in the liver of ACSL4 ^LPC-KO^ and WT mice mice, however, no differences were detected in the investigated cell types.

In order to gain a more precise insight into hepatic inflammation, we further analyzed the cytokines involved in the genesis of MASH in the entire liver tissue, but again the expression of distinct genes was unaffected by ACSL4^LPC-KO^ (see Fig. 3d). Finally, we wished to rule out the possibility that the inhibition of ferroptosis has a direct impact on fibrogenesis, yet by investigating genes involved in the progression of hepatic fibrosis and fat metabolism no differences could be found again (Fig. 3e).

In conclusion, our study shows that the targeted deletion of ACSL4 in LPC in mice fed a CD-HFD has no effect on the formation of a metabolic syndrome, the development of MASLD, or the inflammation and fibrogenesis that are decisive for the progression to liver cirrhosis. The discrepancy between the findings presented in our work and that of Duan et al. remains unclear, warranting a careful consideration of the experimental conditions, such as animal husbandry and the choice of diet used.

## Material and methods

### Mice

*Acsl4*^*tm1a(EUCOMM)Wtsi*^ mice were obtained from Infrafrontier (EMMA strain EM:05887). These were subsequently crossed with Flp-deleter mice to remove the FRT-flanked lacZ/neomycin cassette to obtain mice only harboring the loxP site-flanked Acsl4 alleles (designated *Acsl4*^*tm1c(EUCOMM)Wtsi*^). These mice were then crossed with *alfpAlb*-Cre transgenic mice [36] to generate LPC-specific ablated mice. Only age- and sex-matched floxed littermates were used (WT).

### Diet

In all experiments, 6-week-old male mice were fed with a CD-HFD (Research Diets; D05010402) or with a SD (Ssniff; V1534-30099) for varying time intervals as indicated. At the end of the feeding protocol, mice were fasted at least for 12 h before being euthanized.

### Glucose tolerance test

Mice were fasted overnight. On the next day, 2 mg of glucose per g body weight was intraperitoneally injected and tail blood was taken at the indicated time points. Glucose levels were determined using a hand-held glucose analyzer (Contour XT Bayer) for the indicated time points.

### Serum-Analysis

Serum ALT, AST, GLDH, LDH, cholesterol and triglycerides were measured by standard procedures in the Laboratory Diagnostic Center (LDZ) of the RWTH University Hospital Aachen.

### RNA isolation and cDNA synthesis

RNA isolation and cDNA synthesis were conducted in accordance with the manufacturer’s protocol using the RNeasy Mini Kit (Qiagen) for RNA isolation. The purity and concentration of the RNA were determined spectrophotometrically on the NanoDrop at an absorbance of 260 nm/280 nm. To obtain complementary DNA (cDNA), 1 μg of RNA per experimental sample was processed with the QuantiTect cDNA Synthesis Kit (Qiagen) according to the manufacturer’s instructions. For subsequent processing by real-time polymerase chain reaction (RT-PCR), the cDNA was diluted with nuclease-free water to a concentration of 10 ng/μl.

### Oligonucleotides

The oligonucleotides were purchased from MWG Biotech and were used for quantitative expression analysis by polymerase chain reaction (PCR). According to the manufacturer’s instructions, the oligonucleotides were dissolved in a solution of 100 pg/µl. For practical reaction application, the corresponding primer pairs were diluted again, resulting in a final concentration of 0.4 pg/µl.

### Quantitative real-time polymerase chain reaction

RT-PCR was conducted on the ViiA7 qRT-PCR system (Applied Biosystems/Thermo Fisher Scientific) in 96-well microtiter plates utilizing the oligonucleotide primers previously described. Each reaction batch comprised a total volume of 25 μl, comprising 1.2 μl cDNA (10 ng/μl), 12.5 μl GoTaq qPCR Master Mix (Promega), 9.3 μl nuclease-free H_2_O and 1 μl of each of the corresponding sense and antisense primers (10 pmol/μl). The PCR reaction cycles included a single step of 2 min at 50°C, an initial denaturation at 95°C for 10 min, followed by 40 cycles, each cycle consisting of 15 s at 95°C (denaturation) and 1 min each at 60°C (annealing and elongation). A melting curve analysis was conducted for quality assurance purposes following each run, comprising 15 s at 95°C, 1 min at 60°C, and 15 s at 95°C. For each experimental condition, two ΔCt values were determined by RT-PCR and the arithmetic mean was calculated.

### Western blot Analysis

Liver tissue was homogenized in NP-40 lysis buffer using a tissue grind pestle (Kontes). Cell debris were removed by centrifugation for 10 minutes with 10.000 rpm and 4°C thereby gaining protein lysates. These were resolved by reducing SDS-polyacrylamide gel electrophoresis (PAGE), transferred to PVDF membrane and analyzed by immunoblotting as previously described [37]. Membranes were probed with anti ACSL4 (Invitrogen) and anti GAPDH (Biorad). As secondary antibodies, HRP-conjugated anti-rabbit (ACSL4) and HRP-conjugated anti-mouse (GAPDH)

### Histological examination and evaluation

Liver tissue was fixed with paraformaldehyde (4 %) and paraffin embedded. Resulting paraffin sections (3 µm) were stained with hematoxylin and eosin (H/E) or various primary and secondary antibodies followed by diaminobenzidine (DAB) staining. The following antibodies were used: antibodies against F4/80 (BMA Biomedicals AG, 1:120), B220 (BD Pharmingen, 1:3000) and CD3 (Zytomed, 1:250). The analysis was conducted using QuPath.

The histological scoring system for non-alcoholic fatty liver disease (NAFLD), the old nomenclature of MASDL, was performed according to the NAS scoring system [38].

### Ethics approval

All animal experiments were approved by the Federal Ministry for Nature, Environment and Consumers’ Protection of the state of North Rhine-Westphalia and were performed in accordance to the respective national, federal and institutional regulations. The authors confirm that all experiments were done in accordance to the ARRIVE guidelines.

### Statistical analysis and general experimental design

The data were analyzed using PRISM software (GraphPad Prism 8 Software, Inc., La Jolla, CA) and are expressed as the mean with SD if indicated. The statistical significance between the experimental groups was assessed using an unpaired two-sample t-test and a Mann-Whitney test.

## List of Abbreviations

ACSL4: Acyl-CoA synthetase long-chain family member 4
HCC: Hepatocellular carcinoma
MASLD: Metabolic dysfunction-associated steatotic liver disease
MASH: Metabolic dysfunction-associated steatohepatitis
PCD: Programmed cell death
PUFA: Polyunsaturated fatty acid
MCD: Methionine-choline deficient diet
CD-HFD: Choline-deficient, high-fat diet
WT: Floxed wild-type littermates
LPC: Liver parenchymal cell
HFD: High-fat diet
PC: Phosphatidylcholine
PE: Phosphatidylethanolamine
ROS: Reactive oxygen species
AST: Aspartate aminotransferase
ALT: Alanine aminotransferase
GLDH: Glutamate dehydrogenase
LDH: Lactate dehydrogenase
PCR: Polymerase chain reaction

## Data availability statement

The datasets used and analyzed during the current study available from the corresponding author on reasonable request.

## Author contributions

C.A., M.V. and T. L. designed and guided the research. C.A., M.V. and T. L. wrote the manuscript with help from other authors. C.A. and C.K. performed and analyzed most of the experiments. A.T.S., L.K.., M.T.S, M.C. J.G.B., contributed to research design and/or conducted experiments. M.C. provided the mouse model. All authors read and agreed on the content of the paper.

## Acknowledgments

The authors would like to express their gratitude to Sebastian G. Doll for providing assistance with the ACSL4 mouse model.

## Additional information

The authors declare no competing interests related to this work. M.C. is cofounder and shareholder of ROSCUE Therapeutics GmbH.

